# Comprehensive analyses of the cysteine thiol oxidation of PKM2 reveals the effects of multiple oxidation on cellular oxidative stress response

**DOI:** 10.1101/2020.05.14.097139

**Authors:** Hayato Irokawa, Satoshi Numasaki, Shin Kato, Kenta Iwai, Atsushi Inose-Maruyama, Takumi Ohdate, Gi-Wook Hwang, Takashi Toyama, Toshihiko Watanabe, Shusuke Kuge

## Abstract

Redox regulation of proteins via cysteine residue oxidation is known to be involved in the control of various cellular signal pathways. Pyruvate kinase M2 (PKM2), a rate-limiting enzyme in glycolysis, is critical for the metabolic shift from glycolysis to the pentose phosphate pathway under oxidative stress in cancer cell growth. The PKM2 tetramer acts as pyruvate kinase (PK), whereas the PKM2 dimer, which is induced by Cys358 oxidation, has reduced PK activity. Here, we identified four oxidation-sensitive cysteine residues (Cys152, Cys358, Cys423, and Cys424) responsible for three different oxidation forms. Possibly due to obstruction of the dimer-dimer interface, sulfenylation (-SOH) at Cys424 inhibited tetramer formation and PK activity. Cys423 is responsible for intermolecular disulfide bonds with heterologous proteins. In addition, intramolecular polysulfide linkage (–Sn–, n≧3) possible between Cys152 and Cys358 also is induced. We found that cells expressing the oxidation-resistant, constitutive-tetramer PKM2 (PKM2^C358,424A^) show a higher intracellular reactive oxygen species (ROS) and greater sensitivity to ROS-generating reagents and ROS-inducible anti-cancer drugs. These results highlight the possibility that PKM2 inhibition via Cys358 and Cys424 oxidation contributes to the elimination of excess ROS and oxidative stress.

## Introduction

Oxidative modifications of protein cysteine residues represent post-translational modifications (PTMs). Notably, cysteine (Cys) residue can react with electrophilic-structured compounds such as reactive oxygen species (ROS), resulting in various chemical modifications. Hydrogen peroxide (H_2_O_2_), which is the most abundant ROS in aerobic organisms, reacts to Cys thiol (Cys-SH) and forms various types of oxidized cysteine residues such as disulfide, sulfenic acid, and sulfinic acid (1). Such oxidations alter the function of target proteins and, in many cases, are involved in a signaling cascade for defense mechanisms against oxidative stress (2-7).

Activation of glutaredoxins and peroxiredoxins, which are major antioxidant systems, requires NADPH as a proton donor (8,9). The major pathway for NADPH production is the pentose phosphate pathway (PPP), which is a branch pathway of glycolysis. Therefore, under oxidative stress conditions, increases of the metabolic flux to PPP by glycolysis inhibition serves an important role in the detoxification of H_2_O_2_ (10-12). This metabolic alteration is known to be induced by the ROS-induced inactivation of glyceraldehyde-3-phosphate dehydrogenase (GAPDH) and pyruvate kinase (PK) (13). GAPDH has an oxidation-sensitive (redox-active) cysteine residue in the enzymatic active site. The cysteine residues are evolutionarily conserved and oxidized by various SH-oxidizing reagents and intracellular levels of H_2_O_2_, resulting in many modifications, such as disulfide bond formation (14-20). However, much of the mechanism of PK redox regulation remains unknown. In the present study, we identified multiple redox-active cysteine residues and oxidation forms that affect PK activity.

The PK of a rate-limiting enzyme for glycolysis catalyzes the conversion of phosphoenolpyruvate (PEP) and ADP to pyruvate and ATP (PK activity). This reaction is irreversible and represents the final step in glycolysis. In the yeast model system, inhibition of PK activity induces metabolic change under oxidative stress (21,22). In mammalians, there are four isoforms and two genes, with each isoform expressing in different tissues. The expression of PKL, PKR, and PKM1 is tissue-specific (23), whereas that of PKM2 is ubiquitous. PKM1 and PKM2, encoded by the *PKM* gene, depend on alternative splicing using two mutually exclusive exons (24): PKM1 that includes exon 9 is expressed in limited tissues (such as heart, muscle, brain), whereas PKM2 that includes exon 10 is ubiquitously expressed in many tissues and cells. Interestingly, the regulation of PKM2 enzymatic activity plays a critical role in cancer cell metabolism (25,26). As such, the appropriate regulation of PKM2 might confer metabolic advantages for tumor growth and progression since cancer cells tend to prefer metabolism via glycolysis rather than the oxidative phosphorylation pathway even in aerobic conditions. This is referred to as the Warburg effect (27,28). Tetramer PKM2 and dimer/monomer PKM2 exhibit higher and lower PK activity, respectively. Allosteric activators of PKM2 such as fructose-1,6-bisphosphate (FBP) and serine increase PK activity by promoting the tetramer form (29,30), while the PTMs of PKM2 (e.g., oxidation, phosphorylation, acetylation, and glycosylation) decrease PK activity by blocking the formation of the tetramer. The decrease of PKM2 activity via PTMs provides a metabolic advantage for the Warburg effect and facilitates cancer cell growth (31-34). Thus, the appropriate inhibition of PK activity might induce the accumulation of intermediate metabolite of glycolysis and promote the synthesis of several amino acids and nucleotides including NADPH. As a result, this change fulfills the metabolic requirement for cancer cells (35). Moreover, dimeric PKM2 that is induced by several specific PTMs is responsible for regulation of gene transcription via interacting transcription factors in the nucleus (36,37). Several studies have reported that the moonlighting (non-canonical) functions of PKM2 serve a critical role in tumorigenesis (38). Taken together, in cancer cell growth, the tetramer-to-dimer transition of PKM2 provides two benefits. These are the downregulation of PK activity and activation of moonlighting functions.

Cysteine modifications (oxidations) are important PTMs in PKM2. An initial report indicated that Cys358 oxidation was crucial for oxidative stress-induced downregulation and the intermolecular disulfide bond formation of PKM2 (39). However, the molecular basis of PKM2 oxidation has not been elucidated. Whether another Cys might be responsible for the disulfide bond as a partner has not been resolved. Recently, two reports revealed that cysteine residues at 152, 326, 358, 423, and 424 were involved in modification (40,41). Thus, comprehensive observation of Cys oxidation of PKM2 gives insight into the role of PKM2 in the cellular oxidative-stress response.

## Results

### Identification of novel oxidation-sensitive cysteine residues of PKM2

A previous report indicated that a specific cysteine residue (Cys358) of PKM2 was being involved in its redox regulation using diamide (39). Diamide is a widely utilized thiol oxidant and ROS generator. Recent global analyses on the oxidative modification of protein cysteine residues indicated that another cysteine residue (Cys424) in PKM2 was modified under oxidative stress (42,43). Moreover, as shown in Fig. S1, the PK activity of H1299 cells expressing PKM2 with a loss of Cys358 mutation (PKM2^C358A^) was still downregulated in the presence of H_2_O_2_ (See “**Redox regulation of PKM2 contributes to oxidative stress response***”* section).

Thus, multiple cysteine residues might be responsible for the redox downregulation of PK activity. To identify cysteine residues that might be oxidation-sensitive, we performed a PEG maleimide-based gel shift assay. Sulfhydryl residues of whole-cell proteins were first blocked with N-ethyl maleimide (NEM). Then a reduction of preoxidized Cys via DTT treatment was performed, followed by the addition of PEGlyated maleimide (PEGM, MW 2,333) to NEM-free Cys (NEM-DTT-PEGM assay) (see Fig. 1A). As shown in Fig. 1B, the mobility of PKM2 (63 kDa) in SDS-PAGE shifted to a higher molecular size (100 kDa) when all 10 PKM2 Cys were assumed to be modified by PEGM under the condition without NEM. In contrast, the 63 kDa band was shifted to one band (66 kDa) and two bands (66 kDa and 70 kDa) when HEK293T and HepG2 cells were treated with H_2_O_2_ and diamide, respectively. However, tert-butyl hydroperoxide (tBHP) treatment did not affect the mobility of PKM2 (Fig. 1B). Although this oxidized PKM2 may be specific to the oxidant used, it seems likely that all three oxidation forms were efficiently induced by diamide. We found that H_2_O_2_ level in the diamide-treated cell culture was enhanced (Fig. 1C). Thus, possibly due to the oxidation of cellular reduction systems such as glutathione, diamide induces an intracellular H_2_O_2_ level that might result in more efficient Cys oxidation than the addition of H_2_O_2_ in the culture medium.

**Figure 1.**
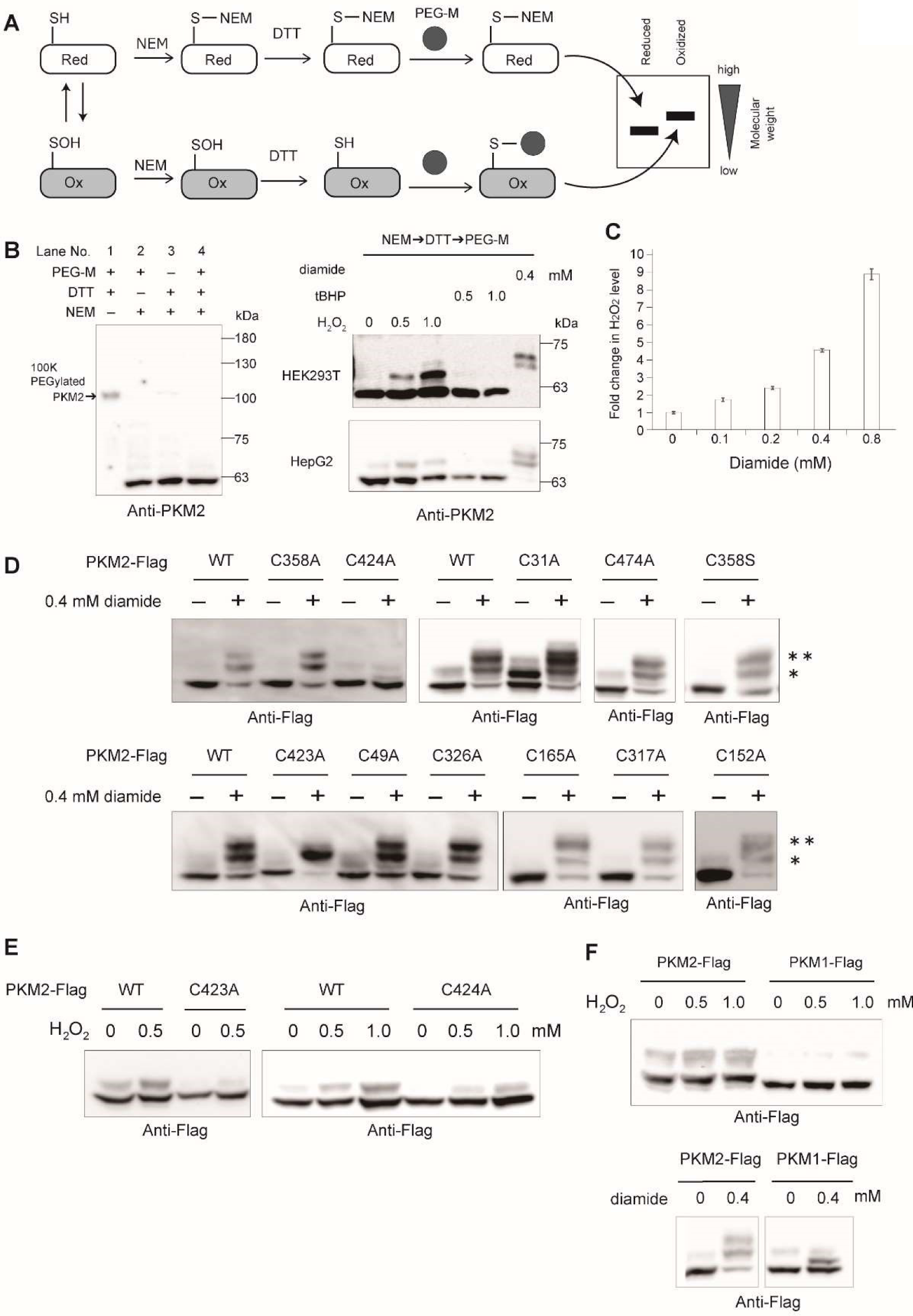
Identification of oxidation-sensitive cysteine residues of PKM2 via NEM-DTT-PEGM assay. **A**. Schematic representation of NEM-DTT-PEGM assay. **B**. Left panel: molecular weight shift of PKM2 by NEM-DTT-PEGM assay in H1299 cells. Each of the three steps was omitted (lanes 1 to 3) and all steps were performed (lane 4) as indicated. Right panels: molecular weight shift (oxidation) of PKM2 in HEK293T and HepG2 cells in response to the indicated concentration (mM) of diamide, tBHT, and H_2_O_2_ in DMEM for 10 min. **C**. Relative levels of H_2_O_2_ in cells’ response to diamide. H1299 cells were treated with the indicated concentration of diamide for 120 min, and the relative H_2_O_2_ level was observed by ROS-Glo H_2_O_2_ assay. Results were normalized to the untreated group. The data are presented as the mean ± the standard error of the mean (N = 3). **D-F.** The oxidation of the PKM2 cysteine mutant was detected by NEM-DTT-PEGM assay in 0.4 mM diamide-treated H1299 shPKM2 cells expressing PKM2^WT^-Flag (WT) or each of the 11 Cys mutants (D, indicated as CxxxA, where xxx is the number of cysteine residues mutated to Ala or as PKM2^CxxxA^-Flag mutants), in 0–1.0 mM H_2_O_2_-treated H1299 shPKM2 cells expressing each PKM2^WT^-Flag, PKM2^C423A^-Flag and PKM2^C424A^-Flag (E), in 0–1.0 mM H_2_O_2_- or 0.4 mM diamide-treated H1299 shPKM2 cells expressing WT PKM2-Flag and PKM1-Flag (F).

Next, to identify cysteine residues in PKM2 responsible for the shift (oxidation), we created 10 PKM2 mutants, in which each of 10 cysteine residues was replaced with alanine (Ala). H1299 shPKM2 cells expressing each of the PKM2 mutants were treated with oxidants. As shown in Fig. 1D, the appearance of the two molecular-weight shifts by diamide were almost completely abolished by Cys424 mutation (PKM2^C424A^), while only the upper-shift band (70 kDa) was absent when Cys423 was mutated (PKM2^C423A^, indicated as ‘**’; Fig. 1D). By contrast, a mutation on Cys358 (PKM2^C358A^ and PKM2^C358S^) did not affect PKM2 oxidation (Fig. 1D). The H_2_O_2_-induced 66 kDa shift was decreased by the mutation of either Cys424 (PKM2^C424A^) or Cys423 (PKM2^C423A^) (Fig. 1E). These results suggest that Cys424 and Cys423—but not Cys358—are found by NEM-DTT-PEGM assay to be sensitive to oxidation induced by H_2_O_2_ and diamide. Notably, Cys424 is a unique Cys mutant present in PKM2 but not in PKM1. Notably, we found that PKM1 was not oxidized by H_2_O_2_, and it was only partially oxidized by diamide (Fig. 1F). Taken together, we identified Cys423 and Cys424 as novel oxidation-sensitive cysteine residues.

### The characterization of oxidative modification of all cysteine residues

The formation of an intramolecular disulfide bond in particular proteins enhances the protein migration in SDS-PAGE under non-reducing conditions (Fig. 2A) (3,44,45). A previous study indicated that faster-migrating PKM2 bands appearing in response to diamide were completely abolished by the mutation of Cys358 to serine (Ser) (39). We observed that the faster-migrating PKM2 appeared in response to 0.4 mM diamide, but not in response to H_2_O_2_ (Fig. 2B). Treatment with DTT (Fig. 2B) and the substitution of Cys358 to Ala— but not Cys424 to Ala—(Fig. 2C) abolished the formation. Moreover, the *in vitro* oxidation assay of recombinant PKM2 proteins (rPKM2) demonstrated that Cys358—not Cys424—is involved in the formation of the faster-migrating band, which is induced by both diamide and H_2_O_2_ (Fig. 2D). Notably, we observed that this faster-migrating band was absent with NEM but not iodoacetamide (IAA) treatment (Fig. 2E). In general, a disulfide bond is not replaced by NEM. Therefore, the DTT-sensitive faster-migrating band of PKM2 may be a NEM-sensitive linkage between Cys358 and other Cys. Recent findings have indicated that polysulfide can link between two cysteine residues (–Sn–, n≧3), and that the linkage is cleaved by NEM but not by IAA (46). Therefore, we investigated the possibility of polysulfide formation using a generator of Cys polysulfur, Na_2_S_4_. The faster-migrating band of PKM2 was strongly induced by Na_2_S_4_, but it disappeared with the addition of DTT and NEM (Fig. 2F). Furthermore, the faster-migrating band induced by Na_2_S_4_ was not detected in the recombinant PKM2^C358A^ mutant (Fig. S2). Based on these results, we concluded that the diamide-induced faster-migrating PKM2 could be due to an intramolecular linkage via polysulfide bond, but not via a disulfide bond between Cys358 and other Cys within the same PKM2 molecule.

**Figure 2.**
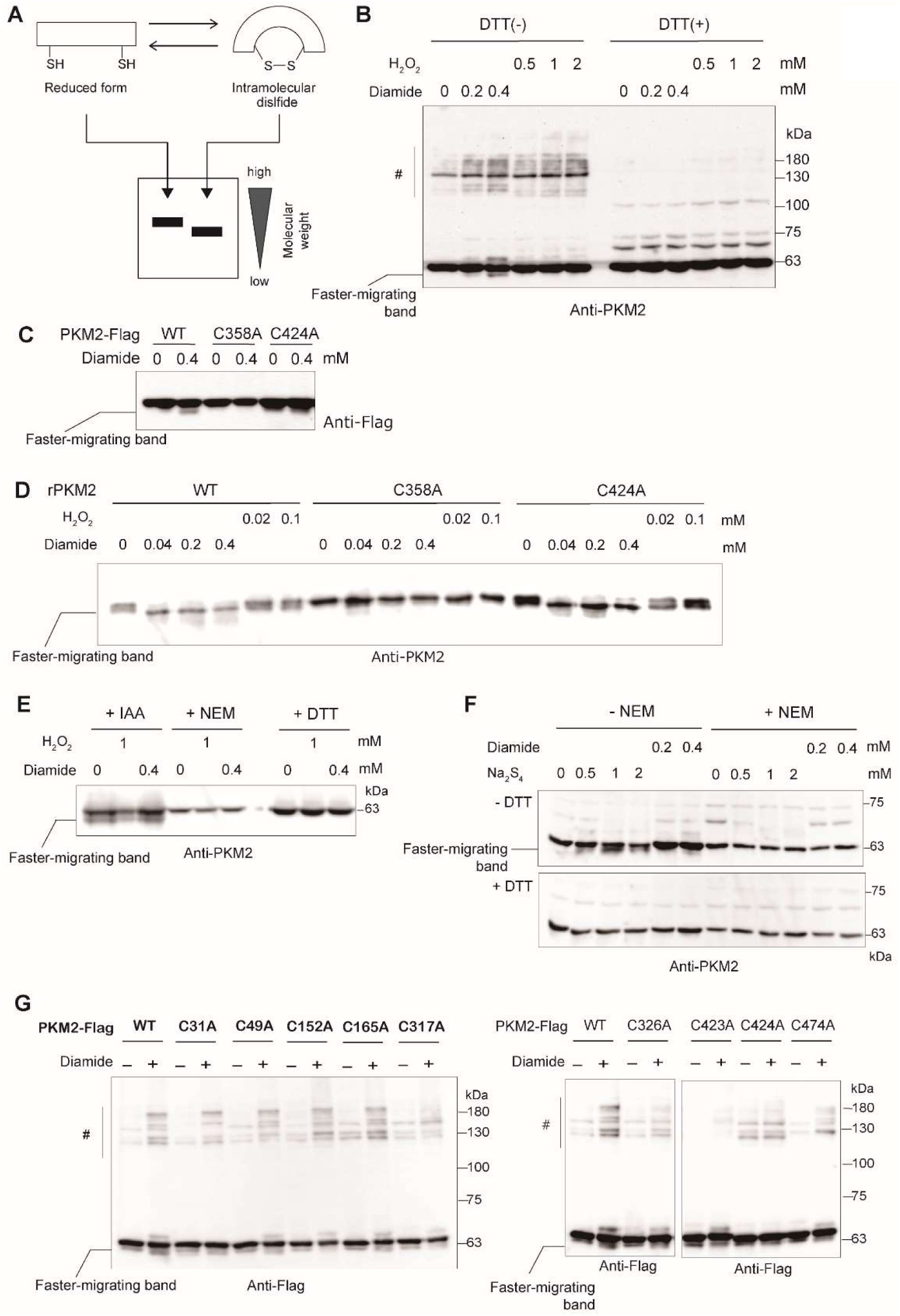
Oxidation of Cys358 is responsible for the possible formation of an intramolecular polysulfide bond. **A**. The formation of an intramolecular bridge enhances mobility of PKM2 in SDS-PAGE. **B.** Detection of DTT-sensitive faster-migrating PKM2 and multiple high-molecular PKM2 bands. H1299 cells were treated with each oxidant (mM) for 10 min in DMEM. For the detection of an intramolecular bridge in PKM2, non-reduced SDS-PAGE (untreated DTT) and reduced SDS-PAGE (treated 50 mM DTT) were performed. Multiple high-molecular PKM2 bands (MW130-180) are indicated as “#”. **C.** H1299 shPKM2 cells expressing PKM2^WT^-Flag and PKM2^CA^-Flag mutants were treated with 0.4 mM diamide for 10 min. Non-reduced SDS-PAGE was performed. **D.** Recombinant proteins of PKM2 (rPKM2) were used for the detection of the faster-migrating band. WT and C358A and C424A of rPKM2 proteins (0.5 µg) were treated with each oxidant (mM) for 10 min in PBS and mixed with a sample buffer to perform non-reduced SDS-PAGE. **E.** The Cys358 linkage in the faster-migrating PKM2 was sensitive to NEM but not IAA treatment. H1299 cells were treated with H_2_O_2_ and diamide for 10 min. Non-reduced SDS-PAGE was performed (+IAA) as described in the Materials and Methods section, except that IAA was replaced with NEM (+NEM). DTT was added in the “-IAA” (+DTT) samples. **F.** H1299 cells were treated with diamide (0.2 and 0.4 mM) or Na_2_S_4_ (0, 0.5, 1, and 2 mM) for 10 min. Non-reduced SDS-PAGE (–DTT) and reduced SDS-PAGE (+DTT) were performed. **G.** H1299 shPKM2 cells expressing PKM2^WT^-Flag or PKM2^CA^-Flag mutants were treated with 0.4 mM diamide for 10 min. Non-reduced SDS-PAGE was performed.

To identify a putative partner Cys for the Cys358-polysulfide linkage, we examined the faster-migrating band formation by using cells expressing each of the Cys mutants of PKM2. As shown in Fig. 2G, the faster-migrating band almost completely disappeared when Cys152 was mutated (PKM2^C152A^ in Fig. 2G). The distance in 3D structure between Cys358 and Cys152 is approximately 36 Å (PDB ID: 4B2D). Therefore, the formation of polysulfide may be required to form the linkage. In addition, we noticed multiple bands with high molecular weight (MW130-180 proteins, as indicated by # in Figs. 2B and 2G) that were sensitive to DTT treatment (Fig. 2B) were induced in response to oxidative stress by both diamide and H_2_O_2_ (Fig. 2B). Since PKM2^C423A^ failed to form a multiple band with high molecular weight (Fig. 2G), Cys423 could be responsible for the intermolecular disulfide bond between PKM2 molecules and/or between a PKM2 molecule and unknown protein(s).

### The sulfenylation of Cys424 decreases the tetramer formation of PKM2

Cysteine-sulfenic acid (Cys-SOH) formation (sulfenylation) is an unstable oxidation form that proceeds to a disulfide bond or Cys-sulfinic acid (Cys-SOOH) (47,48). Recent reports have indicated that protein sulfenylation itself is involved in the redox-dependent regulation of many biological pathways (49,50). Thus, our results suggest that Cys424 is a candidate for the sulfenylation site of PKM2. We used a Cys-SOH reactive compound, dimedone, to detect PKM2 Cys-SOH. Dimedone-reactive protein in PKM2-Flag-immunoprecipitates was detected by the anti-dimedone antibody in PKM2-Flag-expressing cells (Fig. S3). We observed dimedone-reactive PKM2 in cells expressing PKM2^C358A^ and PKM2^C423A^. However, we failed to detect this in cells expressing PKM2^C424A^ and PKM2^C358,424A^ (Fig. 3A). In addition, the sulfenylation of recombinant PKM2 increased in an H_2_O_2_-concentration-dependent manner. However, the rPKM2 sulfenylation was markedly decreased by C424A and C358,424A mutation (Fig. 3B). These results suggest the possibility that Cys424 is a sulfenylation-sensitive Cys in PKM2. Taken together, our results demonstrate that four cysteine residues of PKM2 (Cys152, Cys358, Cys423, and Cys424) are responsible for three different oxidative modifications. These are sulfenylation (Cys424), intramolecular polysulfide formation (Cys358 and Cys152), and intermolecular disulfide formation (Cys423) in PKM2 under oxidative stress (Fig. 3C).

**Figure 3.**
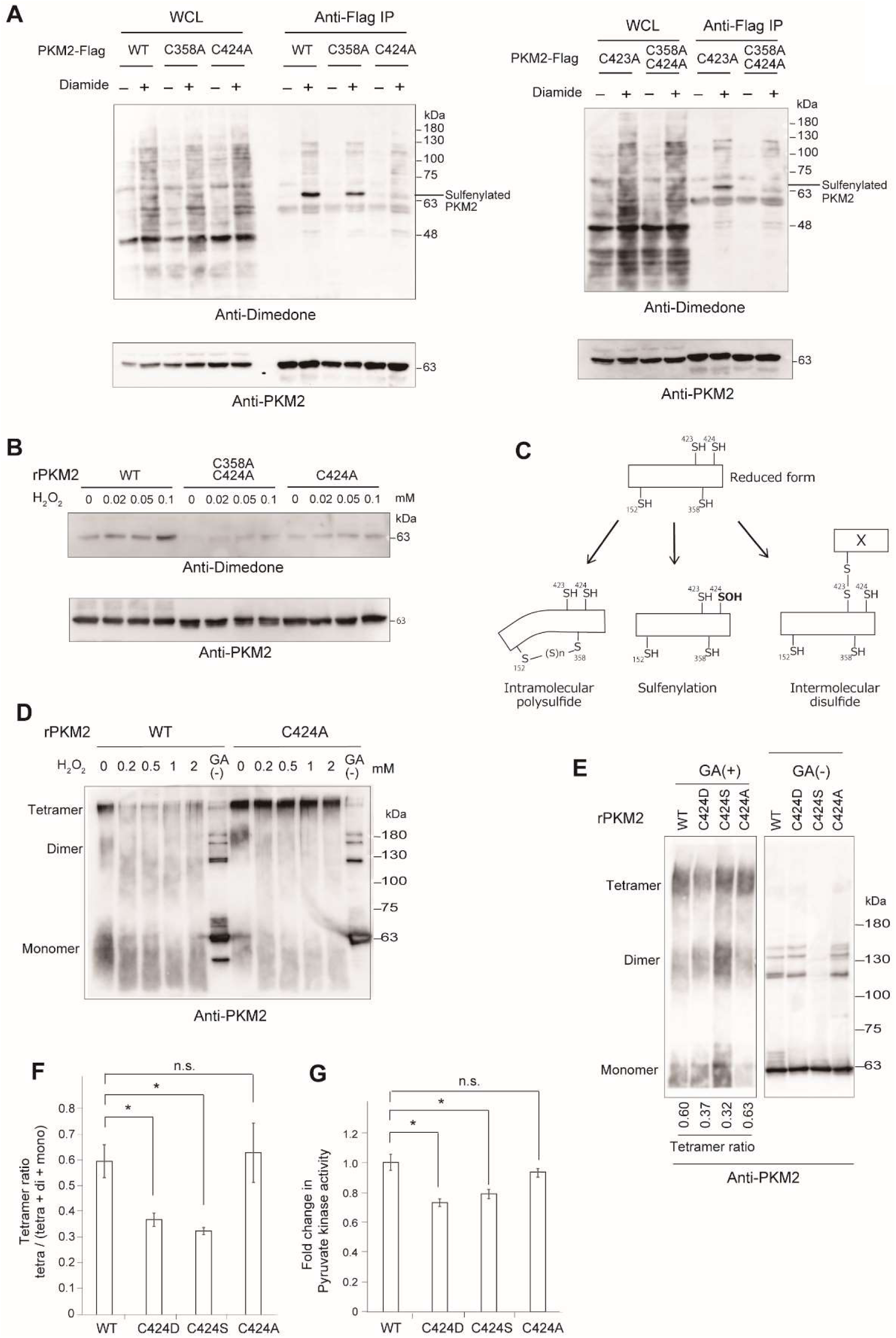
The sulfenylation (-SOH) of PKM2 occurs at Cys424 and inhibits tetramer formation. **A.** Detection of sulfenylated PKM2 in cultured cells. H1299 shPKM2 expressing PKM2^WT^-Flag and PKM2^CA^-Flag mutants were treated with 0.4 mM diamide (+) or without (-) for 10 min and lysed in lysis buffer containing dimedone for the detection of sulfenylation. Immunoprecipitation was performed with the anti-Flag antibody. **B.** Detection of sulfenylated PKM2 in recombinant proteins. rPKM2 WT and rPKM2 Cysteine mutants were treated with H_2_O_2_ and dimedone in PBS for 30 min. **C.** Schematic illustration of cysteine oxidation in PKM2. An open box indicates PKM2, and X indicates an unknown non-PKM2 protein. (D–F) Detection of monomers, dimers, and tetramers of PKM2 *in vitro*. rPKM2 proteins were pre-incubated with FBP and DTT and treated with GA as described in the Materials and Methods section. The non-GA-treated sample is indicated as “GA (-)”. **D.** Inhibition of tetramer formation of rPKM2 by H_2_O_2_ treatment. The indicated concentration of H_2_O_2_ was treated for 120 min before GA treatment. **E.** rPKM2 WT and Cys424 mutants were pre-incubated with DTT and FBP and treated with GA (+) or untreated (-). **F.** The relative intensity of the specific bands of PKM2 tetramer, dimer, and monomer was calculated using Image Lab software (Bio-Rad). The tetramer ratios are expressed as the mean +/- the standard error of the mean (N = 3). Statistical comparisons were performed using Dunnett’s post-hoc test. Differences were considered to be significant at \**p* < 0.05, vs. WT. **G.** rPKM2 WT and rPKM2 Cys424 mutants that were pre-incubated with DTT were diluted with PBS, and the measurement of pyruvate kinase activity was performed. Results were normalized to WT and expressed as the mean +/- the standard error of the mean (N = 3). Statistical comparisons were performed using Dunnett’s post-hoc test. Differences were considered to be significant at \**p* < 0.05, vs. WT.

Next, we investigated the effects of Cys424 oxidation on the multimer formation of PKM2. The PK activity of PKM2 is activated by tetramer formation, whereas dimers and monomers of PKM2 are less active (51,52). We treated cell lysate with GA to crosslink multimer proteins, which are separated by SDS-PAGE under reducing conditions. As previously indicated (53), the tetramer formation of PKM2 was abrogated by S437Y mutation (Fig. S4). We observed that the tetramer was markedly decreased by treatment with diamide (Fig. S4), while the remaining levels of the tetramer under diamide treatment were higher in PKM2 with the simultaneous mutation of C358A and C424A (PKM2^C358,424A^), or even only a C424A mutation (PKM2^C424A^). Thus, the inhibition of tetramer formation may be partly due to the oxidation of Cys424. To clarify the effect of sulfenylation Cys424 on multimer formation, recombinant PKM2s (rPKM2s) were used for further investigation. The tetramer formation of rPKM2^WT^ was significantly decreased by treatment with H_2_O_2_, while its formation of rPKM2^C424A^ was not affected (Fig. 3D). Interestingly, Cys424 is located on the surface of dimer-dimer interaction, and it is considered a crucial residue for tetramer formation (54). Furthermore, a previous study has demonstrated that the mutation of Cys424 to a hydrophobic residue (e.g., leucine, C424L) increases PK activity, whereas, in the case of hydrophilic residues (e.g., serine, C424S) this mutation decreases PK activity (40). As shown in Figs. 3E and 3F, the tetramer ratio of rPKM2^C424A^ is not affected by its mutation (tetramer ratio: WT = 0.6, C424A = 0.63), while the tetramer ratio of the hydrophilic mutations rPKM2^C424D^ (aspartate, C424D) and rPKM2^C424S^, which might mimic Cys424 sulfenylation, were lower than that of rPKM2^WT^ (tetramer ratio: C424D = 0.37, C424S = 0.32, Fig. 3F). Furthermore, the PK activity of these mutants (rPKM2^C424D^ and rPKM2^C424S^) was lower than that of rPKM2^WT^ (Fig. 3G). Taken together, it is possible that sulfenylation of Cys424—which increases the hydrophilicity of Cys-SH—disrupts tetramer formation by increasing hydrophilicity on the dimer-dimer interface and contributes to the inhibition of PK activity.

### Redox regulation of PKM2 contributes to oxidative stress response

To assess the functional significance of the redox regulation of PKM2, we examined the importance of the redox-sensitive cysteine residues on the PK activity and oxidative stress sensitivity of cells. PKM2^WT^-Flag and its Cys mutants were introduced in H1299 cells, of which endogenous PKM2 was stably knocked down using a lentivirus-based shRNA expression vector (H1299 shPKM2 cells; Figs. S5A and S5B). First, we investigated the effect of cysteine oxidation on the PK activity. The aforementioned results indicate that mutations on Cys358 and Cys424 do not affect oxidative stress-induced suppression of PK activity (Fig. S1). Thus, we created a PKM2 mutant with a simultaneous mutation on Cys358 and Cys424 (PKM2^C358,424A^).

The oxidation of PKM2 treated with H_2_O_2_ or diamide was almost completely abrogated in cultured cells expressing PKM2^C358,424A^ (Fig. S6). In addition, the induction of the faster-migrating (the intramolecular polysulfide) rPKM2 by H_2_O_2_ and diamide (Fig. 4A) and the sulfenylated rPKM2 by H_2_O_2_ (Fig. 3B) were not detected in rPKM2^C358,424A^ *in vitro*. Although the PK activity in the lysate of H1299 cells expressing PKM2^WT^ was significantly increased by DTT treatment, the levels of PKM2^C358,424A^ were not increased as mach (Fig. 4B). Similarly, the decreased level of PK activity in the cells expressing PKM2^WT^ in response to H_2_O_2_ and diamide was lower in the cells expressing PKM2^C358,424A^ (Fig. 4B). Thus, these results indicated that Cys358 and Cys424 are responsible for oxidation-induced suppression of PK activity of PKM2.

**Figure 4.**
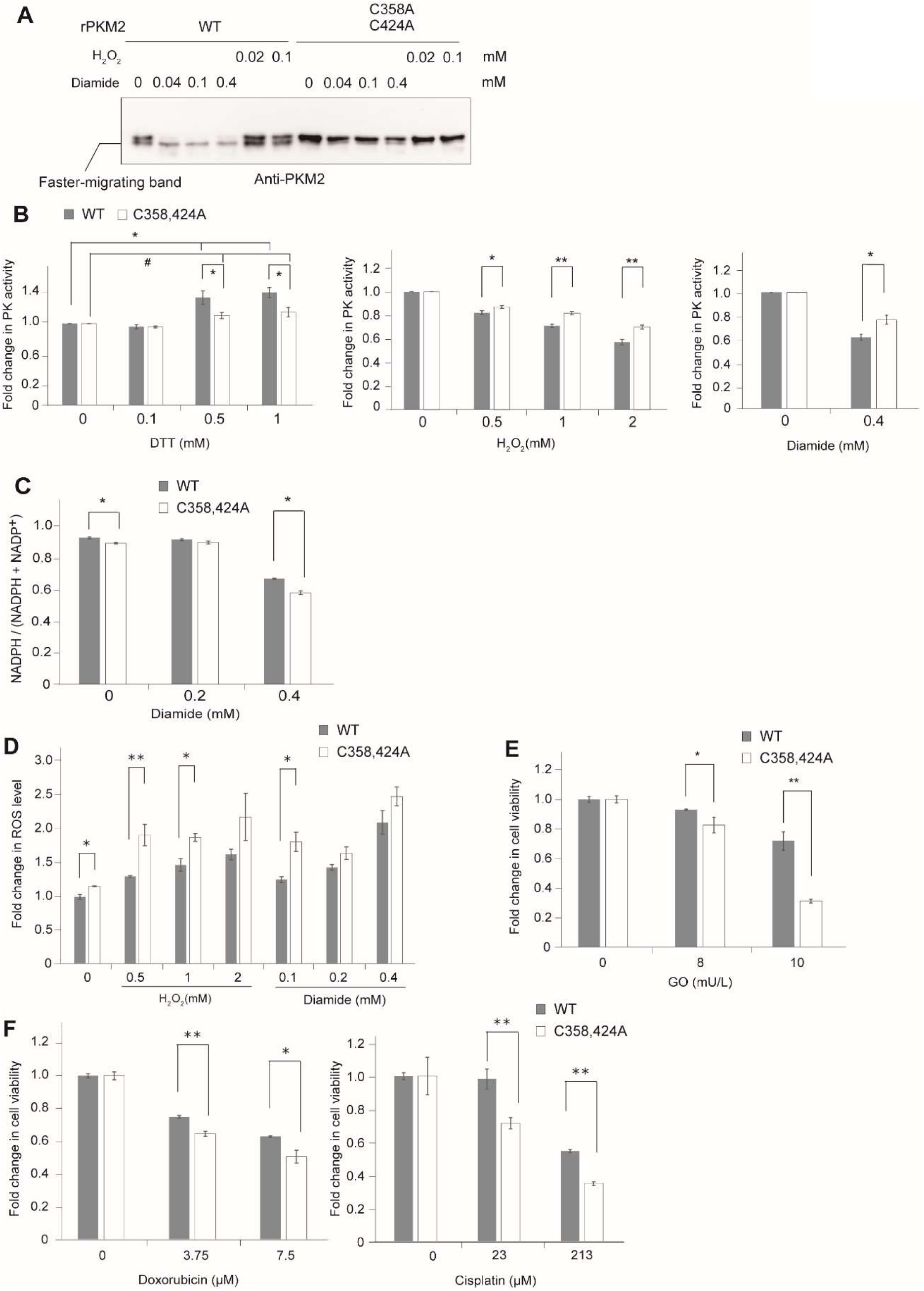
Contribution of PKM2 redox regulation to the oxidative stress response. **A.** rPKM2^WT^ and the rPKM2^C358,424A^ mutant were treated with each oxidant (mM) for 10 min. Non-reduced SDS-PAGE was performed to detect an intramolecular bridge of PKM2. **B.** Effect of C358,424A mutation on PK activity in cells under reduction and oxidation conditions. H1299 shPKM2 cells expressing PKM2^WT^-Flag (closed bar) and those expressing PKM2^C358,424A^-Flag mutant (open bar) were treated with the indicated concentration of H_2_O_2_ and diamide for 10 min, respectively. The cells were then lysed in lysis buffer and PK activity was observed. For DTT experiments, lysate prepared from cells without oxidant treatment was treated with the indicated concentration of DTT 37°C for 10 min before activity was measured. **C.** H1299 shPKM2 cells expressing PKM2^WT^-Flag and PKM2^C358,424A^-Flag mutants were treated with diamide for 30 min. The NADPH ratio was observed as described in the Materials and Methods section. **D.** H1299 shPKM2 cells expressing PKM2^WT^-Flag (closed bar) and PKM2^C358,424A^-Flag mutant (open bar) were treated with each oxidant for 30 min. To measure intracellular ROS, we used CellRox Orange Reagent as described in the Materials and Methods section. The fluorescence intensity was normalized to WT without oxidant treatment. **E, F.** H1299 shPKM2 cells expressing PKM2^WT^-Flag (closed bar) and PKM2^C358,424A^-Flag mutant (open bar) were treated with glucose oxidase (GO) or anti-cancer drugs overnight. Cell viability was determined by Alamar blue assay (Thermo Fisher Scientific) according to the manufacturer’s protocol. Results were normalized to the untreated group. (B–F) The data are presented as the mean +/- the standard error of the mean (N = 3). Statistical comparisons were performed using the Student’s t-test. Differences were considered to be significant at * *p* < 0.05, ** *p* < 0.01, and to be not significant at # *p* > 0.05.

The inhibition of PKM2 enzymatic activity alters the flow of metabolites into PPP, and it contributes to NADPH generation and oxidative stress response (39,55). Therefore, we investigated the intracellular level of NADPH and the accumulation of intracellular ROS under oxidative stress conditions. The NADPH ratios (NADPH/(NADPH+NADP)) in cells were decreased in response to the diamide treatment (Fig. 4C). These decreases were marked in cells expressing PKM2^C358,424A^ (Fig. 4C). In addition, the intracellular ROS levels were enhanced in response to oxidative stress (H_2_O_2_ and diamide). Again, ROS levels in response to H_2_O_2_ and diamide in cells expressing PKM2^C358,424A^ were higher than in those expressing PKM2^WT^ (Fig. 4D). Next, to examine the effect of continuous oxidative stress on cell viability, we added glucose oxidase (GO) to the culture medium. This generates H_2_O_2_ by glucose catalysis (56). Cells expressing PKM2^C358,424A^ were more sensitive than those expressing PKM2^WT^ (Fig. 4E). These results highlight the possibility that PKM2 inhibition via Cys358 and Cys424 oxidation contributes to the elimination of excess ROS and oxidative stress.

Some anti-cancer drugs induce intracellular ROS, which gives rise to cytotoxic cancer cells (57,58). Therefore, the ability to eliminate ROS is an important factor for chemosensitivity. We tested whether the oxidation of PKM2 is involved against the chemosensitivity of cisplatin (CDDP) and doxorubicin (DOX). As shown in Fig. 4F, the cells expressing PKM2^C358,424A^ were more sensitive than those expressing PKM2^WT^. These results suggest that the oxidative stress response via the Cys358 and Cys424 oxidation in PKM2 might be involved in the sensitivity of CDDP and DOX. Collectively, our findings indicate a mechanism for oxidative stress resistance via the downregulation of PKM2 by redox-based modification of Cys358 and Cys424 to enhance NADPH levels through the potential activation of PPP under oxidative stress conditions.

## Discussion

In the present study, we demonstrated the existence of three different oxidation statuses of PKM2 (sulfenylation, disulfide, polysulfide). First, we identified Cys423 and Cys424 as novel redox-active cysteines using NEM-DTT-PEGM assays (Figs. 1A, 1B, and 1D). This method facilitates the detection of NEM-non-reactive yet DTT-sensitive cysteine oxidation, which includes Cys-sulfenylation (SOH) and Cys-Cys disulfide. Our results suggest that oxidation induces Cys424 sulfenylation (Figs. 3A and 3B). Cys424 is located at the interface of the dimer-dimer interaction of the tetramer PKM2. Interestingly, Cys424 is a unique cysteine residue in PKM2, but not in PKM1. The corresponding residue of PKM1 is leucine. It has been shown that increasing hydrophobicity at the 424 residues (e.g., from C424A to C424L) enhanced PKM2 in an active state (tetramer) because of increased dimer-dimer surface interaction (54). Conversely, Asp and Ser substitution of PKM2 Cys424, which might decrease hydrophobicity, suppresses the formation of the active form (Figs. 3E and 3F). Thus, Cys424 sulfenylation might increase hydrophilicity on the dimer-dimer interface, thereby leading to decreased tetramer formation (Figs. 3D–3F), suppression of PK activity (Fig. 3G), and oxidative stress resistance (Fig. 4E). Second, we also demonstrated that PKM2 Cys423 is involved in intermolecular disulfide bonding with other proteins (Fig. 2G). However, since the PKM2^C423A^ mutant was enzymatically inactive (Fig. S7), we could not pursue further analysis to understand the effect of Cys423 oxidation on PK activity. Since the corresponding cysteine residue in PKM1 was conserved, the possible partner protein linking Cys423 may be crucial for PK activity and important for identification in future studies. Third, we revealed that Cys358 and Cys152 are essential for the formation of a faster-migrating band under non-reduced SDS-PAGE (Fig. 2C and 2G). The band completely disappeared following treatment with an SH-alkylation reagent, NEM (Figs. 2E and 2F). An intramolecular disulfide bond, which is induced by H_2_O_2_ in many cases, is not reactive to NEM. Recently, Ida et al. have reported protein S-polythiolation, which is more unstable and highly reactive (59). We observed that an intracellular generator of reactive-sulfur species (RSS) induced intramolecular Cys-Cys linkage (Fig. 2F). This suggests that an intramolecular polysulfide linkage between Cys152 and Cys358 enhances the faster-migrating band. Although diamide induced possible Cys S-polythiolation of PKM2, it was unknown whether intracellular RSS was induced by diamide, and its oxidation mechanism remains unclear.

The inhibition of PKM2 by oxidants contributes to the metabolic change required to enhance metabolites in PPP (39). Our findings indicated that cells expressing the oxidation-resistant form of PKM2 (PKM2^C358,424A^) increased intracellular ROS, and they were more sensitive to ROS-generating reagents, such as glucose oxidase, and ROS-inducible anti-cancer drugs, such as cisplatin and doxorubicin (Figs. 4D–4F). Thus, the inhibition of PKM2 via Cys358 and Cys424 oxidation is important to the oxidative stress response.

In the tumor progression process, cancer cells are exposed to higher levels of oxidative stress (60). As such, compounds that increase intracellular ROS are considered anti-cancer agents that enhance cell death (61,62). PK activity may be repressed due to the oxidation of PKM2 in the oxidative tumor environment. Our findings suggest that the specific inhibition of Cys358 and Cys424 oxidation to reduce PK suppression may represent ideal targets for anti-cancer drugs.

## Experimental procedures

### Cell lines, cell culture, and the establishment of PKM2 knockdown

A human non-small cell lung carcinoma cell line H1299 and HepG2 were obtained from American Type Culture Collection (ATCC). H1299 is a human non-small cell lung carcinoma cell line. HepG2 is a hepatocellular carcinoma HepG2 cell line. HEK293T (Human Embryonic Kidney cells 293) cells were obtained from Thermo Fisher Scientific, MA, USA. These cells were cultured in DMEM (Nissui, Tokyo, Japan or Nacalai Tesque, Kyoto, Japan) supplemented with 10% fetal bovine serum (Biosera, Kansas, MO, USA), 3% glutamine (Nacalai Tesque), penicillin/streptomycin (FUJIFILM Wako, Osaka, Japan), and 7.5% NaHCO_3_ (Nacalai Tesque) at 37°C and 5% CO_2_ concentration.

Stable knockdown of endogenous PKM2 was performed with a lentivirus expressing a high pyruvate kinase knockdown efficiency shRNA (63). The shPKM2 sequence was cloned into AgeI-EcoRI sites of a pLKO.1 puro vector (pLKO-shPKM). The lentivirus vectors were produced in HEK293FT cells by the co-transfection of plasmids pMD2.G and pCMVR8.74 with the pLKO.1-sh PKM2 using Lipofectamine 2000 (Thermo Fisher Scientific, Waltham, MA, USA). To produce H1299 shPKM2 cells, H1299 cells were transduced by the lentivirus vectors and selected with DMEM containing 5 µg/ml puromycin (InvivoGen, San Diego, CA, USA).

pMD2.G (Addgene plasmid # 12259; http://n2t.net/addgene:12259; RRID: Addgene_12259) and pCMVR8.74 (Addgene plasmid # 22036; http://n2t.net/addgene:22036; RRID: Addgene_22036) were provided by Didier Trono. pLKO.1 neo was provided by Sheila Stewart (Addgene plasmid # 13425; http://n2t.net/addgene: 13425; RRID: Addgene_13425).

### Expression of PKM2 in H1299 shPKM2 cells

Human PKM2 cDNA (NM_002654.6) was fused with a Flag-tag to the corresponding C-terminus of the PKM2 sequences. PKM2-Flag was cloned into a pEB multi-hygro vector (FUJIFILM Wako) between *Kpn*I and *Not*I. The PKM2 coding sequence corresponding codon 106-112 was modified as 5’-GCcGTcGCcCTg-3’ (the lowercased letters indicate substitutions) to perform shRNA-resistant expression. To constract PKM2 cysteine mutants, corresponding cysteine codon was mutated to Ala (GCC) or Asp (GAC) or Ser (AGC) using PCR. These plasmids were then transfected in H1299 shPKM2 cells using FuGENE HD transfection reagent (Promega, Madison, WI, USA) and cultured in DMEM for 48 h (Fig. S5A).

### Antibodies and reagents

The primary antibodies used in this study were rabbit anti-PKM2 (SAB4200095, Merck, Darmstadt, Germany), mouse anti-Flag M2 (F1804, Merck), rabbit anti-SOH (07-2139, Merck), mouse anti-GAPDH (Merck), and rabbit anti-actin (Santa Cruz, Dallas, TX, USA). Secondary antibodies included polyclonal goat anti-rabbit HRP (Dako, Agilent, Santa Clara, CA, USA) and polyclonal goat anti-mouse IgG HRP (Dako, Agilent). The oxidants used in this study were 30% hydrogen peroxide (H_2_O_2_, Nacalai Tesque), diamide (MP Biomedicals, Irvine, CA, USA), tBHP (Merck), and glucose oxidase (GO, Merck). The cysteine residue modification reagents used were N-ethyl maleimide NEM (Nacalai Tesque), iodoacetamide IAA (FUJIFILM Wako), PEGylated maleimide PEGM (MW 2,000) (SUNBRIGHT®, ME-020MA, NOF Corporation, Tokyo, Japan), and dimedone (Nacalai Tesque). DTT (Nacalai Tesque) was used for the reduction of oxidized cysteine residues. Additional chemical reagents used were sodium tetrasulfide Na_2_S_4_ (Dojindo, Kumamoto, Japan), cisplatin CDDP (FUJIFILM Wako), and doxorubicin DOX (FUJIFILM Wako).

### Sodium dodecyl sulfate-polyacrylamide gel electrophoresis (SDS-PAGE) and western blotting

The proteins were quantified with a DC Protein Assay Kit (Bio-Rad, Hercules, CA, USA) using BSA (Nacalai Tesque) as the standard. The proteins prepared in each experiment were mixed SDS sample buffer (50 mM Tris-HCl, pH 6.8, 2% SDS, 0.1% bromophenol blue, 10% glycerol) supplemented with 50 mM DTT and boiled at 95°C for 5 min. We performed the SDS-PAGE in running buffer (25 mM Tris-HCl, pH 7.5, 250 mM Glycine, 0.1% SDS). Proteins in the gel were then transferred to a poly-1,1-difluoroethane (PVDF) membrane (Merck) in transfer buffer (25 mM Tris-HCl, pH 7.5, 250 mM glycine, 0.1% SDS, 5% methanol) using a trans-blot system (Bio-Rad). Blocking the PVDF membrane was performed by shaking in TBS-T (10 mM Tris-HCl, pH 7.5, 150 mM NaCl, 0.1% Tween 20) containing 5% skim milk for 1 hr at room temperature. After the primary- and secondary-antibody reactions, we obtained chemiluminescent images using Immobilon Western Chemiluminescent HRP Substrate (Merck) and the VersaDoc imaging system (Bio-Rad) or the ChemDoc imaging system (Bio-Rad).

### Maleimide-based gel shift assay (NEM-DTT-PEGM assay)

After cells were treated with each oxidant at the indicated time, they were incubated for 5 min in 100 mM NEM in PBS (Nacalai Tesque) to block free thiols. Cells were then lysed in PK lysis buffer (50 mM Tris-HCl, pH 7.5, 1 mM EDTA, 150 mM NaCl, 1% Igepal-630 (Merck)) supplemented with 50 mM DTT and protease inhibitors (cOmplete™ protease inhibitor cocktail EDTA-free, Roche, Basel, Switzerland)). Lysates were centrifuged (13,800 g, 5 min, 4°C). Supernatants were collected in new tubes and then incubated for 10 min at room temperature to reduce the oxidized thiols. To remove DTT, protein in these lysates were precipitated in 5% trichloroacetic acid (TCA)-75% acetone for 10 min on ice. The proteins were precipitated by centrifugation (13,800 g, 2 min, 4°C). The pellets were then washed with 500 µL acetone and collected by centrifugation. This step was repeated three times to neutralize the pellets. After neutralization, the pellets were suspended again in urea PEGM buffer (50 mM Tris-HCl, pH 7.5, 150 mM NaCl, 8 M urea, 1% SDS, 5 mM PEGM). These samples were then mixed in SDS sample buffer (50 mM Tris-HCl, pH 6.8, 10% glycerol, 2% SDS, 0.01% bromophenol blue) supplemented with 50 mM DTT. SDS-PAGE was then performed within 30 min of sample preparation.

### Non-reduced SDS-PAGE

After cells were treated with each oxidant for the indicated time, they were incubated for 5 min in 100 mM IAA in PBS to block free thiols. Cells were then lysed in PK lysis buffer supplemented with 50 mM IAA and protease inhibitors. Lysates were centrifuged (13,800 g, 5 min, 4°C), and the supernatants were collected in new tubes. These samples were mixed with SDS sample buffer (without DTT), and SDS-PAGE was performed.

### Purification of recombinant PKM2 (rPKM2)

Recombinant PKM2 (rPKM2) WT—or its mutants fused with a His-tag on the amino-terminus—were expressed in *Escherichia coli* BL21 (DE3) using pET15b vector (Merck). A single colony was inoculated in 2 mL Luria-Bertani broth (LB) supplemented with ampicillin (LB Amp) and was incubated overnight at 37°C with shaking. Overnight cultures were diluted in 400 mL LB Amp and grown to an OD_600_ of 0.4–0.6 at 37°C with shaking. Then, 1 mM isopropyl-β-D-thiogalactopyranoside (Nacalai Tesque) was added to these cultures to induce the expression of His-PKM2, followed by incubation at 25°C for 6 hr with shaking. Cells were harvested by centrifugation (4,000 g, 4°C, 10 min) and pellets were stored at -20°C.

The pellets were lysed in 10 mL Buffer A (PBS, 20% glycerol, 10 mM imidazole, 0.1% Tween 20, 1 mg/ml lysozyme chloride) via sonication using Vibra-Cell™ (VC505, Sonics & Materials, CT, USA). After the lysates were centrifuged, the supernatants were mixed with Ni-NTA (Qiagen, Germantown, MD, USA), which is pre-equilibrated by Buffer A before use with rotation at 3 hr in 4°C in Econo-Pac Chromatography Columns (20 mL, Bio-Rad). After the unbounded fraction was discarded, Ni-NTA was washed twice with 4 mL Buffer B (PBS, 20% Glycerol, 20 mM imidazole, 0.1% Tween 20). His-tagged PKM2 protein was eluted by Buffer C (PBS, 20% Glycerol, 250 mM imidazole, 0.1% Tween 20) and dialyzed at 4°C overnight in dialysis buffer (50 mM Tris-HCl, pH 7.5, 100 mM KCl, 10 mM MgCl_2_, 20% glycerol, 20% ammonium sulfate, 0.05% Tween 20). The purified proteins were checked and quantified using a Coomassie brilliant blue (CBB) staining (Quick-CBB, FUJIFILM Wako) with BSA as the standard (Fig. S8).

### Detection of PKM2 sulfenylation in cultured cells

H1299 shPKM2 cells transfected with pEB PKM2-Flag were treated with oxidants. To block free thiols, cells were incubated with 100 mM NEM in PBS at 37°C. Then, cells were washed with PBS and lysed in a dimedone-containing lysis buffer (50 mM Tris-HCl, pH 7.5, 150 mM NaCl, 5 mM dimedone (Nacalai Tesque), 1% Igepal, protease inhibitors). After removal of the unresolved fraction by centrifugation, the supernatants as whole-cell lysates (WCLs) were quantified using a DC Protein Assay kit (Bio-Rad). Then, 600 µg WCL was mixed with anti-Flag antibody beads (Merck) at 4°C for 3 hr with rotation. The beads were then washed with the lysis buffer three times. Bound proteins were eluted from the beads with 90 µL of sample buffer containing 50 mM DTT at 95°C for 5 min. The immunoprecipitates were analyzed by SDS-PAGE and western blotting using the anti-PKM2 antibody and rabbit anti-SOH (anti-dimedone) antibody, which specifically reacts with dimedone-bound cysteine residues.

### Detection of rPKM2 sulfenylation

rPKM2 WT or its mutants (0.5 µg) were diluted in PBS. These proteins were treated with H_2_O_2_ for 10 min. Then, dimedone was added to the protein sample for a final concentration of 0.25 mM and incubated at room temperature for 30 min. The reaction mixtures were combined with a sample buffer supplemented with 2-mercaptoethanol (Nacalai Tesque) and analyzed by SDS-PAGE and western blotting, as previously described.

### Detection of rPKM2 multimer

rPKM2 WT or its mutants (0.5 µg) were diluted in PBS and pre-incubated with 1 mM FBP and 5 mM DTT at 37°C for 30 min. Glutaraldehyde (GA) (Nacalai Tesque) was added to the protein sample for a final concentration of 0.025% and incubated at 37°C for 3 min. The reaction mixtures were then mixed with a sample buffer supplemented with 50 mM DTT and analyzed by SDS-PAGE and western blotting using the anti-PKM2 antibody.

### Detection of PKM2 multimer in cultured cell

Cells were lysed in PBS with the addition of 1% Igepal, and protease inhibitors. GA was added to the cell lysate for a final concentration of 0.025% and incubated at 37°C for 3 min. The reaction mixtures were then mixed with a sample buffer supplemented with 50 mM DTT and analyzed by SDS-PAGE and western blotting using the anti-Flag antibody.

### Measurement of pyruvate kinase activity

PK activity was measured by monitoring the change in absorbance at 340 nm because of the oxidation of NADH. This was a coupled reaction with lactate dehydrogenase (LDH).

Recombinant proteins (0.12 µg) and cell lysates (20 µg) were diluted in PK reaction buffer (50 mM Tris-HCl, pH 7.5, 100 mM KCl, 5 mM MgCl_2_). The final reaction mixture was created by adding 1.9 mM PEP (FUJIFILM Wako), 0.6 mM ADP (Merck), 0.3 mM NADH (Merck), 1.1 units LDH (Merck), 1 mM FBP (Merck) (where applicable), and DTT (where applicable). The final reaction volume was 150 µl in a 96-well plate (Greiner, Austria), and the change at 340 nm was measured by VariosKan Flash (Thermo Fisher Scientific).

### Measurement of NADPH ratio and intracellular ROS

Measurement of NADPH ratio was performed using an NADP/NADPH-Glo™ assay kit (Promega), according to the manufacturer’s protocol. We used CellRox Orange Reagent (Thermo Fisher Scientific) and ROS-Glo (Promega) to measure total H_2_O_2_ level in the cell culture (medium and cells). For CellRox Orange, cells were incubated with oxidants at the indicated time in a 96-well plate (Perkin Elmer, Waltham, MT, USA). After removing the medium and washing with PBS, the cells were stained by adding CellRox orange and Hoechst (Thermo Fisher Scientific) at 37°C for 30 min in the dark. The cells were then washed with PBS three times. Fluorescence imaging analysis was subsequently performed using the Operetta CLS High-content Analysis System (Perkin Elmer).

For ROS-Glo, the cells were incubated in 384 white plates (Thermo Fisher Scientific), and the measurement of ROS was performed according to the manufacturer’s protocol (Promega).

## Acknowledgments

We thank Daisuke Sugawara, Satomi Sasaki, and Yukie Sato (Tohoku Medical and Pharmaceutical University) for their technical assistance.

## Author Contributions

Most of the experiments were performed by H. Irokawa. H. Irokawa and S. Kuge designed the research and analyzed the data. S. Numasaki and S. Kato performed the experiments presented in Figure 1 and contributed analytical tools. K. Iwai, T. Ohdate and Inose-Maruyama performed plasmid cloning and contributed to the establishment of the PKM2 knockdown and expression of PKM2-Flag cells. T. Toyama, Gi-W. Hwang, and T. Watanabe consulted and advised on the research. S. Kuge involved in conducting all experiments and wrote the paper.

## Footnotes

## Abbreviation used

Ala: alanine
CDDP: cisplatin
Cys: cysteine
DOX: doxorubicin
FBP: fructose-1,6-bisphosphate
GA: glutaraldehyde
GAPDH: glyceraldehyde-3-phosphate dehydrogenase
GO: glucose oxidase
IAA: iodoacetamide
LB: Luria-Bertani broth
NEM: N-ethyl maleimide
PEGM: PEF maleimide
PEP: phosphoenolpyruvate
PK: pyruvate kinase
PKM2: pyruvate kinase M2
PPP: pentose phosphate pathway
PTMs: post-transcriptional modifications
PVDF: poly-1,1-difluoroethane
ROS: reactive oxygen species
Ser: serine
tBHP: tert-butyl hydroperoxide
WCL: whole cell lysate

